# How Much Coordination Does an AAA+ Motor Require? A Case Study of the Archaeal Cdc48/p97 Homologue

**DOI:** 10.64898/2026.11.20.768179

**Authors:** Paige Kanters, Rui Huang

## Abstract

AAA+ (ATPases Associated with diverse cellular Activities) motors harness ATP hydrolysis to remodel substrates, often through a spiral staircase architecture which is commonly associated with a sequential ‘hand-over-hand’ translocation mechanism. However, the extent to which motor activity requires strict sequential order and interprotomer coordination remains incompletely understood. Here, we investigate the functional resilience of an archaeal AAA+ unfoldase, VAT, using reconstituted heterohexamers containing wild-type protomers and variants defective in nucleotide binding, hydrolysis, substrate engagement, or regulatory control. Quantitative modeling of steady-state ATPase activity and substrate unfolding reveals motif-dependent partial cooperativity. ATP hydrolysis is best described by local interprotomer coupling, whereas substrate unfolding exhibits a threshold-dependent response to defects in conserved mechanochemical motifs. Walker A and Arginine-finger lesions inhibit neighboring ATPase activity but are partially tolerated during unfolding. Pore-loop mutations stimulate ATP turnover while more strongly impairing unfolding. In contrast, N-terminal domain-mediated autoinhibition follows an independent model in both assays, consistent with controlling catalytic output predominantly at the individual protomer level while preserving interprotomer coordination for function. Together, these findings reveal that ATP turnover and substrate unfolding respond differently to local perturbations, with tolerance to protomer defects determined by the mechanochemical element affected. This motif- dependent resilience highlights the operational plasticity of an AAA+ ring and challenges strictly sequential models in which a single defective protomer arrests the motor.

**SIGNIFICANCE:** AAA+ molecular motors are commonly thought to operate through sequential ATP hydrolysis and coordinated conformational changes around a hexameric ring. However, whether such strict coordination is essential for motor function remains unclear. By systematically introducing defective protomers into the archaeal unfoldase VAT, we demonstrate that these motors exhibit remarkable, motif-dependent functional resilience. Defects in ATP binding and hydrolysis are partially tolerated, whereas disruption of substrate-engaging pore loops more severely compromises unfolding despite stimulating ATP turnover. In contrast, regulatory autoinhibition primarily affects individual protomers without imposing comparable constraints on its ATP hydrolysis and unfolding functions. These findings demonstrate that ATP hydrolysis and global mechanical unfolding are separable properties. Our results challenge rigid sequential models and reveal how AAA+ motors balance local mechanochemical coupling with the flexibility required to sustain productive substrate processing.

## INTRODUCTION

AAA+ (ATPases Associated with diverse cellular Activities) proteins are a super family of molecular motors that harness ATP binding and hydrolysis to remodel proteins or other macromolecular substrates. They participate in diverse cellular processes, including protein quality control, membrane remodeling, organelle biogenesis, DNA replication, and gene regulation.^1–3^ Many AAA+ unfoldases and translocases assemble into hexameric rings, in which nucleotide-dependent conformational changes are coupled to substrate engagement and movement through a central pore.^1–3^ High-resolution cryo-electron microscopy (cryo-EM) structures of substrate-engaged AAA+ motors have revealed a recurring architecture that features an asymmetric, right-handed spiral staircase in which conserved axial pore loops contact the polypeptide backbone of the substrate, while one or more subunits disengage at the seam of the ring.^2,4–8^ These structures have motivated a sequential ‘hand-over-hand’ model, in which sequential ATP hydrolysis around the ring drives directional substrate translocation.^2,4,6,9^ However, it is unclear whether these structural snapshots imply an obligatory reaction sequence or represent one accessible transition state in a more complex and flexible mechanochemical cycle.^1,10,11^ Indeed, biochemical, kinetic, and single-molecule studies reveal considerable mechanistic plasticity and diversity in the actions of AAA+ motors.^12–16^ ClpX, for example, retains substrate unfolding and translocation activities with only a subset of catalytically competent subunits, supporting a probabilistic mechanism that does not require strictly sequential hydrolysis.^12,17–19^ Studies of ClpB indicate that substrate unfolding and translocation occur on different timescales,^15^ while recent single-molecule fluorescence studies reveal that Hsp104 displays biased stochastic transitions between inter-subunit conformations and bidirectional substrate movement.^13,16^ Together, these observations challenge an obligatory rigid sequence of conformational changes to facilitate motor function in AAA+ proteins and suggest that they can accommodate structural and kinetic constraints through potentially alternative routes of action. Yet, how such mechanistic plasticity emerges from the interactions between individual protomers remains unclear.

We address this question using VAT, a homohexameric type II AAA+ ATPase from *Thermoplasma acidophilum* and an archaeal homologue of Cdc48/p97. VAT comprises an autoinhibitory N-terminal domain (NTD) and two tandem ATPase domains, D1 and D2 (Figure 1A).^5,20^ High-resolution cryo-EM structures show that it adopts the spiral-staircase architecture in the substrate-engaged state (Figure 1B),^4^ characteristic of many AAA+ motors, making it a representative system for investigating how ATP hydrolyzing, substrate-gripping, and regulatory elements contribute to local coupling and collective unfolding. Specifically, do nucleotide-binding or catalytic defects compromise the entire ring, or can the motor accommodate impaired protomers? Do pore-loop lesions impose different constraints, and can regulatory domains suppress ATP turnover without disrupting collective substrate processing? Comparing these perturbations provides a quantitative means to distinguish requirements for chemical turnover from those for productive mechanical work.

**Figure 1.**
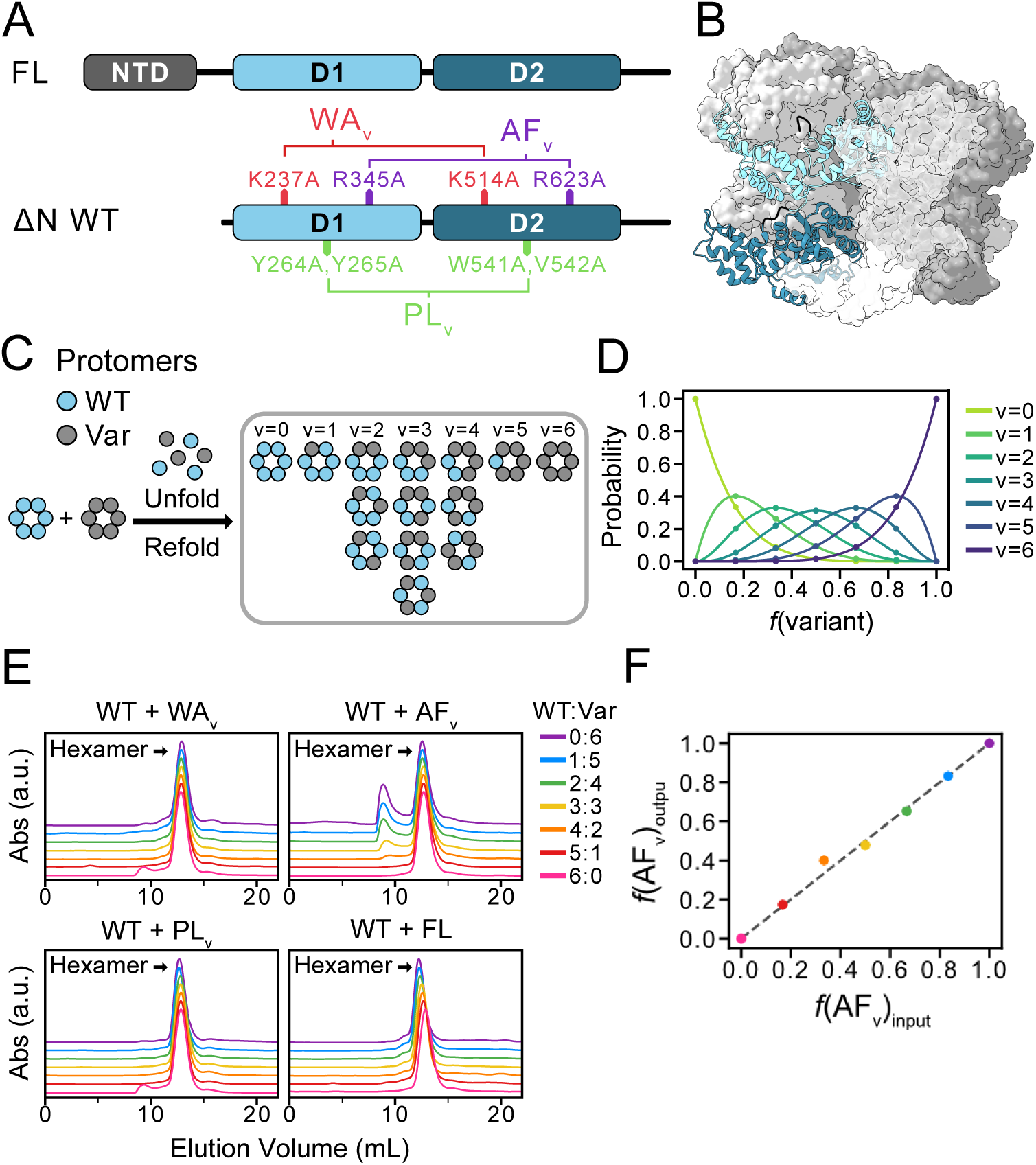
Reconstitution and validation of VAT heterohexamers. (A) Domain organization of full- length VAT (FL) and ΔN-VAT (WT), indicating Walker A (WA_v_), Arginine-finger (AF_v_) and pore loop (PL_v_) mutations. (B) Cryo-EM structure of substrate-engaged and ATPγS-bound ΔN-VAT (PDB ID: 5VCA),^4^ showing the spiral staircase architecture of the hexameric ring. (C) Unfolding–refolding strategy, yielding 14 distinct ring configurations. (D) Predicted populations of heterohexamers containing *v* variant protomers based on the initial mixing ratio and binomial distribution. (E) SEC chromatograms of refolded heterohexamers obtained from different initial mixing ratios. (F) Validation of protomer composition in the mixed rings containing WT and AF_v_ protomers. Corrected relative ESI-MS peak intensities of the refolded samples (Figure S2) are plotted against the input mixing ratios. The grey dashed line indicates y=x.

## MATERIALS AND METHODS

### Reconstitution of Heterohexamers

To reconstitute mixed heterohexamers containing WT and variant protomers (WA_v_, AF_v_, PL_v_, FL), purified WT and variant homohexamers were mixed at the desired molar ratios to achieve a final total protomer concentration of 15 µM in unfolding buffer [50 mM HEPES (pH 7.5), 100 mM NaCl, 1 mM EDTA, 6 M guanidine hydrochloride]. The protein mixtures were then refolded by rapid dropwise dilution (1:24, v/v) into refolding buffer [50 mM HEPES (pH 7.5), 100 mM NaCl, and 1 mM EDTA]. The refolded samples were concentrated and purified by size exclusion chromatography (SEC) to isolate correctly assembled hexamers, which eluted at approximately 12.5 mL. Fractions corresponding to the hexamer peak were pooled, concentrated, flash-frozen in liquid nitrogen, and stored at -80 °C.

### NADH-Coupled ATPase Assay

Steady-state ATPase activity of mixed VAT heterohexamers was measured using an NADH-coupled ATPase assay. Reaction mixtures were prepared in a 96-well plate to a final volume of 100 µL per well. The final reaction mixtures contained 25 mM HEPES (pH 7.5), 100 mM NaCl, 5 mM MgCl_2_, 257 μM NADH, 2 mM ATP, 1 mM phosphoenolpyruvate (PEP), 5.4 U/mL lactate dehydrogenase (LDH), and 2.03 U/mL pyruvate kinase (PK). The final VAT protomer concentrations were 1 µM for WT+WA_v_, WT+AF_v_, and WT+FL mixtures, and 0.25 or 0.15 µM for WT+PL_v_ mixtures. ATP hydrolysis was monitored by measuring the decrease in NADH absorbance at 340 nm using an Epoch 2 microplate reader (Agilent BioTek) at 20 °C. All measurements were performed in triplicate, and the ATPase activity was determined from the linear portion of the absorbance-versus-time traces.

ATP hydrolysis rates were averaged across technical triplicates for each sample and normalized to the activity of the WT homohexamer measured within the same biological replicate (*A*_WT_ = 1). The normalized activities were then averaged across biological replicates, and the standard deviation was calculated. The mean normalized ATPase activity was plotted against the fraction of variant protomers and fit to the proposed cooperativity models (SI Material and Methods).

### Substrate Unfolding Assay

Substrate unfolding activity of mixed VAT heterohexamers was measured using a photoconvertible fluorescent model substrate comprising mEos3.2 fused to an unstructured N-terminal His_6_-tag. Prior to the unfolding assay, the mEos3.2 substrate was irradiated with 365-nm UV light for 3 hours at 4 °C using a 3UV transilluminator (UVP) to induce photoconversion from green to red fluorescence.

For the substrate unfolding assay, reaction mixtures were prepared in a 96-well plate to a final volume of 100 µL per well. The final reaction mixtures contained 25 mM HEPES (pH 7.5), 100 mM NaCl, 5 mM MgCl_2_, 2 mM ATP, 10 mM phosphoenolpyruvate (PEP), and 4.06 U/mL pyruvate kinase (PK). The final protein concentrations were 10 µM VAT protomer and 2 µM photoconverted mEos3.2 substrate. Green fluorescence (excitation/emission: 480/516 nm) and red fluorescence (excitation/emission: 540/580 nm) were monitored using a Synergy H1 microplate reader (Agilent BioTek) at 20 °C. All measurements were performed in triplicate, and unfolding activity was determined from the initial rate of decrease in red fluorescence.

Substrate unfolding rates were averaged across technical triplicates for each sample and normalized to the activity of the WT homohexamer measured within the same biological replicate (*A*_WT_ = 1). The normalized activities were then averaged across biological replicates, and the standard deviation was calculated. The mean normalized unfolding activity was plotted against the fraction of variant protomers and fit to the proposed cooperativity models (SI Material and Methods).

Additional information concerning cloning, protein expression, and purification, as well as the mechanistic models is provided in SI Materials and Methods.

## RESULTS AND DISCUSSION

### Reconstitution and Validation of Compositionally Defined VAT Heterohexamers

To elucidate inter-protomer coordination, we reconstituted mixed heterohexamers from wild-type ΔN- VAT (WT) and variants that carry mutations at key mechanochemical motifs, including the Walker A nucleotide-binding motif (WA_v_), the catalytic Arginine finger (AF_v_), and the substrate-engaging pore loop (PL_v_), as well as the full-length VAT (FL) which contains an autoinhibitory NTD (Figure 1A; Figure S1). Heterohexamers were assembled using an established denaturing–refolding protocol with defined initial mixing ratios between WT and a variant (Figure 1C).^4,14,21–23^ Random incorporation was expected to yield binomially distributed compositions spanning 14 distinct ring configurations (Figure 1C,D).^14,22^ Size- exclusion chromatography (SEC) of the refolded mixtures showed a predominant peak at 12.5 mL, consistent with the formation of hexameric assemblies (Figure 1E). Notably, while AF_v_-containing samples also displayed a minor fraction of soluble aggregates (Figure 1E, ∼ 9 mL), electrospray ionization mass spectrometry (ESI-MS) of the isolated hexamer fraction confirmed protomer compositions consistent with the input mixing ratios (Figure 1F; Figure S2). Further SDS-PAGE analysis provided complementary support for conserved protomer compositions of the WT/FL assemblies (Figure S3), establishing a basis for comparing functional assays across mixed rings with defined compositional distributions.

### Motif-Dependent Interprotomer Coupling Governs ATP Hydrolysis

We first measured steady-state ATP hydrolysis across the compositional titrations. Each titration samples a distribution of 14 discrete ring configurations with binomially defined populations (Figure 1C,D); therefore, the measured bulk rate represents a population-weighted average across all configurations (Figure S4). To quantify interprotomer coupling during ATP hydrolysis, we evaluated four models spanning a range of cooperative behaviors—Independent, All-or-None, Unidirectional, and Bidirectional (Figure 2A). The Independent model assumes the activities are additive and unaffected by their neighbors (Figure 2A, Independent). At the opposite extreme, the All-or-None model assumes that a single defective or hyperactive protomer abolishes or enhances activity across the entire ring (Figure 2A, All-or-None). Because ATP hydrolysis occurs at discrete sites formed at protomer interfaces, we propose two partial cooperativity models assuming local coupling between neighboring protomers. In these two models, the activity of a given protomer, *i*, is allosterically modulated either by its clockwise neighbor (Unidirectional) or by both flanking neighbors (Bidirectional) (Figure 2A). The coupling parameters, α (Unidirectional) and α′ (Bidirectional), describe the magnitude and direction of the neighboring effect, with values near zero indicating little coupling, negative values indicating inhibition, and positive values indicating stimulation (Figure 2B,C). Models were evaluated and compared using reduced chi-squared (***χ***^2^) and corrected Akaike information criteria (AICc) (Figure 2D; Table S1).

**Figure 2.**
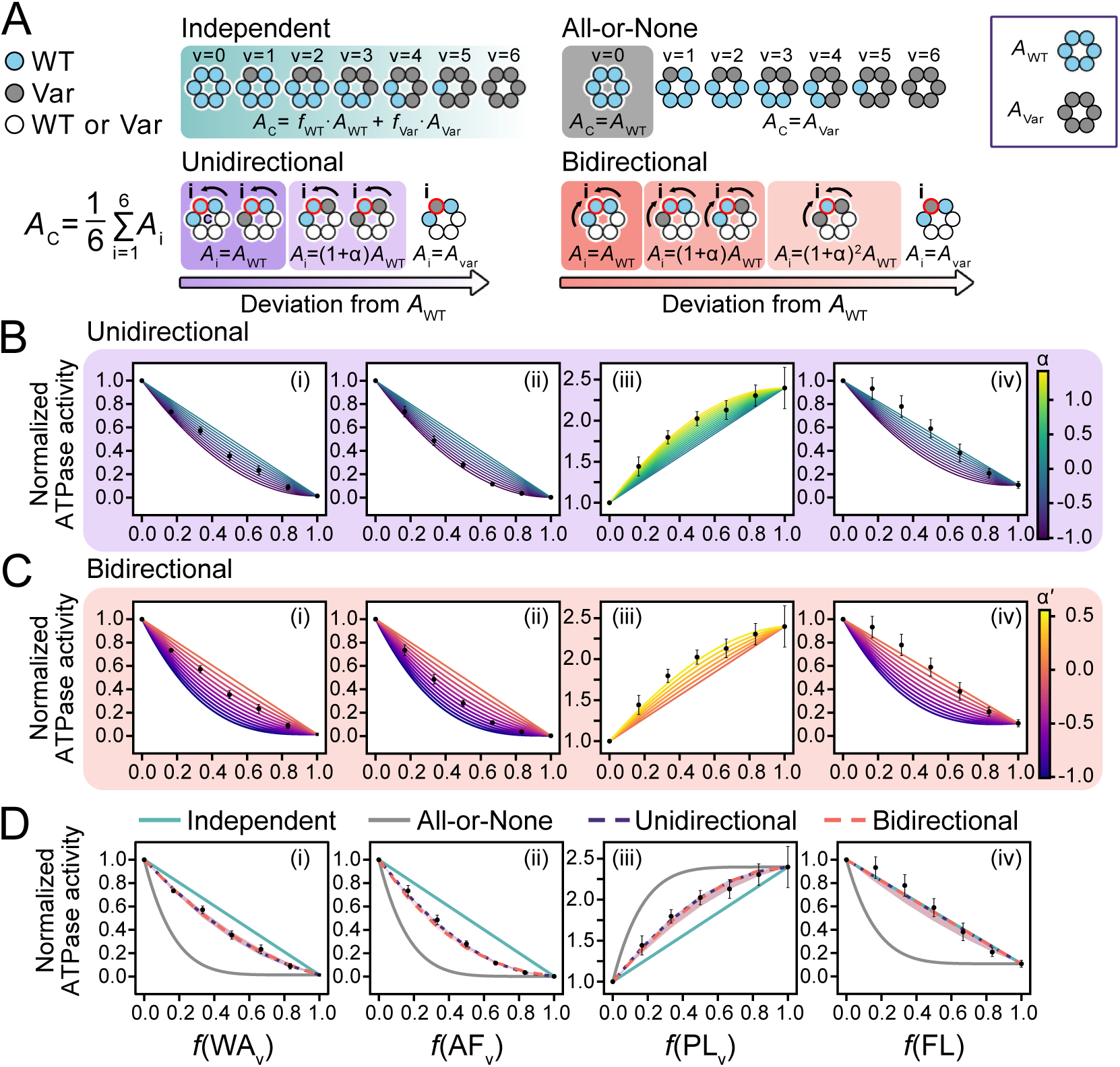
Inter-protomer coupling during ATP hydrolysis. (A) Schematics of the Independent, All-or- None, Unidirectional, and Bidirectional models, relating the ATPase activity of the ring, *A*_C_, to ring configuration. In the Independent model, the activity of the ring is the composition-weighted mean of the activities of WT and variant homohexamers (*A*_WT_ and *A_Var_*). In the All-or-None model, incorporation of one or more variant protomers (v > 0) shifts the activity of the hexamer to that of the variant homohexamer. In the Unidirectional model, the activity of each protomer *A*_i_ depends on the identity of its clockwise neighbor, whereas in the Bidirectional model, it depends on the identities of both adjacent neighbors. In both Unidirectional and Bidirectional models, the activity of the ring, *A*_C,_ is the mean activity of the six protomers. (B, C) Normalized ATPase activities of reconstituted mixed hexamers containing WT and variant protomers, including WA_v_ (i), AF_v_ (ii), PL_v_ (iii), and FL (iv). Experimental data (black circles) are plotted against fractions of the variant protomers and compared with simulated curves for the Unidirectional (B) and Bidirectional (C) models across different values of the coupling parameter, ***α*** or ***α***^#^, respectively. (D) Normalized ATPase activities of the same mixed hexamers, shown with predictions from the Independent (solid teal), All-or-None (solid gray), best-fit Unidirectional (dashed purple), and best-fit Bidirectional (dashed orange) models. Shaded regions indicate 95% confidence intervals (CIs) of the model fits.

Our results revealed that inter-protomer coupling during ATP hydrolysis varies across functional motifs (Figure 2B-D). For WA_v_-, AF_v_-, and PL_v_-containing rings, neither the Independent (Figure 2D, solid teal) nor the All-or-None (Figure 2D, solid gray) model adequately captured the observed activity profiles, supporting partial cooperativity and inter-protomer allostery. Incorporation of either WA_v_ or AF_v_ protomers produced inhibitory effects extending beyond the loss of activity associated with the mutated site, consistent with suppression of neighboring WT protomers (*α*, *α’* < 0) (Figure 2B,C, i,ii). For WA_v_-containing rings, the Unidirectional and Bidirectional models described the data comparably well (***χ***^2^ = 0.71 and 0.60; ΔAIC_C_ = 0.56 and 0.00, respectively). For AF_v_-containing rings, the Unidirectional model was favored (ΔAICc = 0.00 *versus* ΔAICc= 4.41 for Bidirectional), supporting an asymmetric model of neighboring contribution. Using the Unidirectional model for comparison between the effect of WA_v_ and AF_v_, AF_v_ exerts a stronger inhibitory effect on the neighboring protomer (*α* = −0.97 ± 0.02) than WA_v_ (*α* = −0.62 ± 0.05) (Figure 2D, i,ii, dashed purple). This difference indicates that disrupting transition-state stabilization through the Arginine finger and impairing nucleotide binding through the Walker A motif do not have equivalent effects on neighboring ATPase sites.

In contrast, the PL_v_ variant produced the opposite response in ATP hydrolysis. Homohexameric PL_v_ exhibited an ATP turnover rate 2.40 ± 0.25-fold that of WT, highlighting the functional communication between substrate-engaging pore loops and the ATPase machinery.^2,20,24,25^ In mixed rings, PL_v_ also displayed partial cooperativity with the neighbors (Figure 2B,C, iii), but PL_v_ stimulates the ATPase activity of neighboring WT protomers rather than inhibiting. The fitted coupling parameter (*α* = +1.29 ± 0.10, Unidirectional) corresponds to more than a doubling of modeled activity for the neighboring WT protomer (Figure 2D, iii, dashed purple). Thus, pore-loop mutations stimulate ATP turnover in both the mutant protomers and the WT neighbors.

The autoinhibitory NTD displays a distinct inter-protomer effect. Homohexameric full-length VAT (FL) hydrolyzed ATP at 0.11 ± 0.03 of the rate measured for ΔN-VAT (WT), consistent with strong autoinhibition. However, FL-containing rings were best described by the Independent model, rather than the partial-cooperativity models that captured WA_v_, AF_v_, and PL_v_ behavior (Figure 2D, iv). This result is consistent with NTD-mediated inhibition acting predominantly within individual protomers, with little detectable effect on the ATPase activity of neighboring protomers. The NTD therefore appears to regulate catalytic output through a functional response distinct from lesions in nucleotide-binding, catalytic, or substrate-engaging elements.

### Substrate Unfolding Exhibits Motif-Dependent Resilience and Threshold Cooperativity

We next examined how these perturbations affect substrate unfolding using the fluorescent protein mEos3.2 as a model substrate (Figure S5). Unlike ATP turnover occurring at individual protomer interfaces, unfolding requires productive substrate engagement and coordinated action within the central pore shared by all the protomers, suggesting that its dependence on ring composition may differ from that of ATP hydrolysis. Therefore, alongside the Independent and All-or-None models, representing the limiting cases of no cooperativity and complete ring-level dependence, we evaluated three additional models to describe partial cooperativity (Figure 3A). In the Threshold model, the ring retains WT unfolding activity with increasing variant protomers until the number of variant protomers reaches a threshold, *T*, at which activity shifts to the variant level (Figure 3A,B). The Exponential model describes a progressive decrease in activity with each additional variant protomer, governed by the parameter β which ranges from 0 (no inhibition) to 1 (complete inhibition) (Figure 3A,C). The combined Threshold-Exponential (T- E) model allows activity to decline progressively before the threshold *T*, beyond which it assumes the variant activity (Figure 3A,D). These models provide a quantitative means of comparing functional tolerance of substrate unfolding to defects in different structural elements.

**Figure 3.**
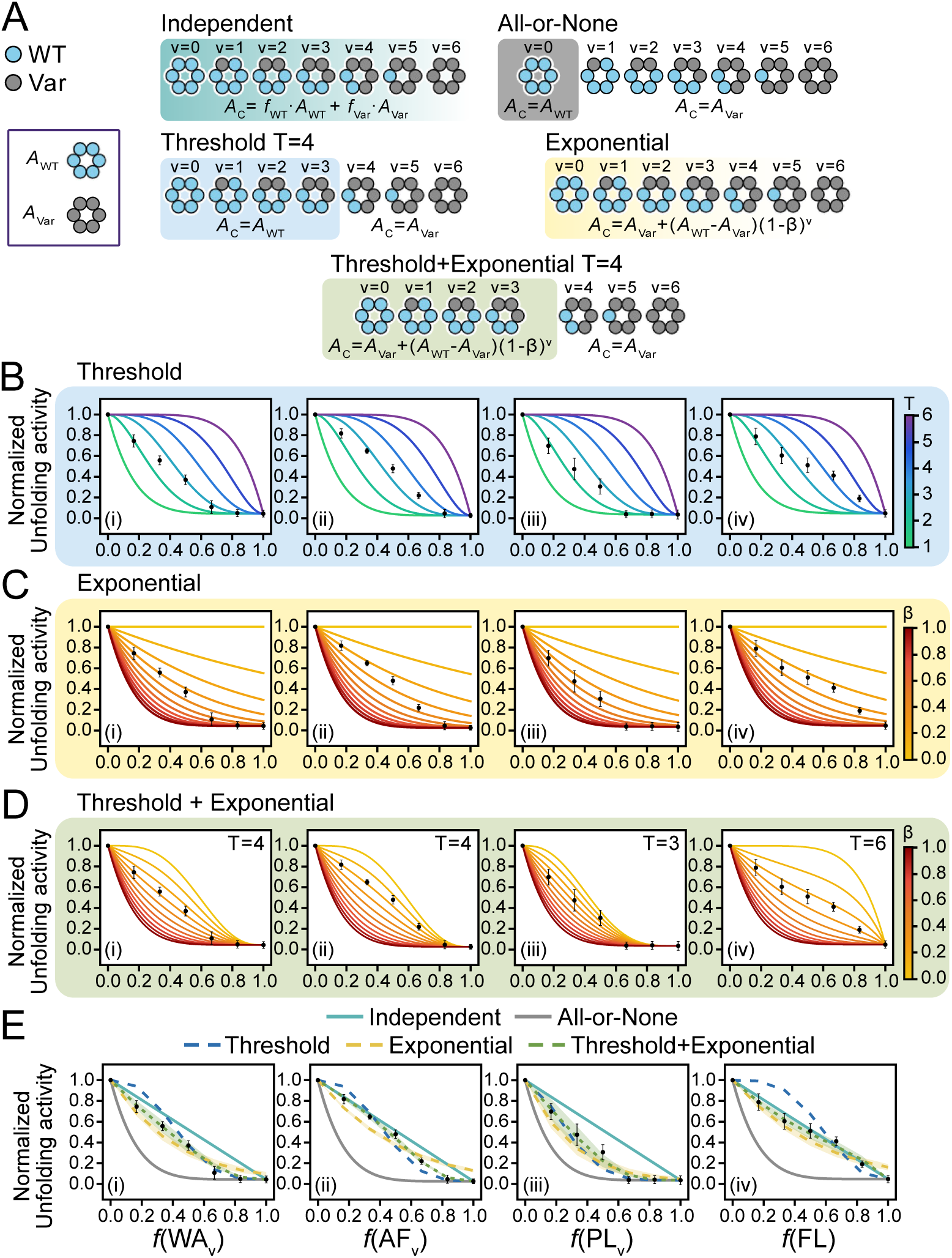
Inter-protomer dependence of substrate unfolding. (A) Schematics of the Independent, All- or-None, Threshold, Exponential, and Threshold–Exponential (T-E) models relating the unfolding activity of the ring, *A*_C_, to ring configuration. Independent and All-or-None models are defined in the same way as in Figure 2A. In the Threshold model, activity is maintained at the WT level until the number of variant protomers reaches a threshold, *T*, beyond which it shifts to the variant level. In the Exponential model, activity is exponentially penalized with each additional variant protomer, with the severity governed by the parameter *β*. In the T-E model, activity is exponentially penalized with increasing variant protomers until the threshold, *T*, is reached, at which point it shifts to the variant homohexamer level. (B-D) Normalized unfolding activities of reconstituted mixed hexamers containing WT and variant protomers, including WA_v_ (i), AF_v_ (ii), PL_v_ (iii), and FL (iv). Experimental data (black circles) are plotted against fractions of the variant protomers, alongside simulated curves for the Threshold (B), Exponential (C), and T-E (D) models across different values of the model parameters, *T* and *β*. (E) Normalized unfolding activities of the same mixed hexamers, shown with predictions from the Independent (solid teal), All-or- None (solid gray), best-fit Threshold (dashed blue), best-fit Exponential (dashed yellow), and best-fit T-E (dashed green) models. Shaded regions indicate 95% confidence intervals (CIs) of the model fits.

WA_v_-, AF_v_-, and PL_v_-containing rings all showed composition-dependent unfolding behavior that departed from both Independent and All-or-None predictions, indicating that unfolding is neither simply additive nor abolished by one defective protomer. The nature of the partial cooperativity, however, depended on the mutated motif. WAv- and AF_v_-containing rings were best described by the T-E model (Figure 3E, i,ii, dashed green; Table S2). Grid searches of ***χ*^2^** identified a threshold of *T* = 4 for both variants (Figure 3D, i,ii; Figure S6), indicating that unfolding could be sustained with up to three variant protomers under the conditions tested based on this model. Below this threshold, the initial penalty was greater for WA_v_ (*β* = 0.28 ± 0.02) than for AF_v_ (*β* = 0.18 ± 0.01). Thus, while both perturbations compromise the ATPase cycle, their effects on unfolding are not identical. Although AF_v_ more strongly inhibited the ATPase activity of the neighbors, WA_v_ imposed more severe unfolding penalty below the shared threshold. These distinct responses show that the effect of a catalytic lesion on bulk ATP turnover does not directly predict its effect on substrate unfolding.

In comparison, PL_v_-containing rings were more sensitive to the incorporation of defective protomers. The Threshold model (Figure 3B, iii) with *T* = 2 and the T-E model with *T* = 3 (Figure 3D, iii) described the data equally well (Figure 3E, iii, dashed blue *versus* dashed green), with the latter implying a substantial initial progressive penalty (*β* = 0.31 ± 0.07). Although these fits did not resolve a unique threshold for PL_v_, both fits indicate that substrate unfolding has a lower tolerance to disruption in the pore loops compared to WA_v_ and AF_v_. This sensitivity underscores the importance of maintaining continuous and productive contact with the polypeptide substrate during unfolding. Moreover, stimulated ATP turnover in PL_v_ and its neighboring WT protomers does not lead to enhanced substrate unfolding, representing uncoupling of chemical turnover from mechanical output, as previously observed for VAT and ClpX.^20,24,25^ Collectively, the results show that perturbations to the ATPase cycle and disruptions to the substrate-gripping element have distinct effects on substrate unfolding.

Finally, FL-containing rings were best described by the Independent model for their unfolding activity, mirroring their behavior in ATP hydrolysis (Figure 2D, iv). These rings retained unfolding activity even when only one or two ΔN WT protomers were present (Figure 3E, iv), in striking contrast to WA_v_-, AF_v_- or PL_v_-containing rings, indicating that NTD-bearing protomers are more readily accommodated than variants with defective mechanochemical motifs. The combined ATPase and unfolding results support that NTD-mediated autoinhibition acts predominantly at the level of individual protomers while remaining compatible with collective substrate processing. Regulatory suppression, therefore, need not impose the same functional constraints as damages to the core mechanochemical machinery. This distinction is highly consistent with the physiological role of the NTD, which regulates catalytic output while still permitting the assembled motor to process protein substrates.

## CONCLUSION

In summary, systematic reconstitution of mixed VAT hexamers with defective protomers or protomers with regulatory moieties allowed us to dissect the operational plasticity and resilience of the AAA+ ring. Our results reveal that although interprotomer cooperativity is essential for efficient ATP turnover and substrate unfolding, the AAA+ ring of VAT can tolerate defective protomers and retain partial ATPase and unfoldase activity in a motif-dependent manner. Walker A and Arginine-finger lesions produce inhibitory interprotomer coupling in ATP hydrolysis but can be partially tolerated during substrate unfolding. Pore-loop mutations instead stimulate ATP turnover while imposing a more stringent constraint on productive unfolding, consistent with uncoupling of chemical turnover from mechanical output. Distinct from defects within the core mechanochemical machinery, NTD-mediated autoinhibition is described predominantly by an Independent model, highlighting that the regulation largely occurs at the protomer level while preserving interprotomer coordination within the ring to sustain function. Together, these findings support a model in which local catalytic regulation and collective substrate processing are related but separable properties of the AAA+ ring, and the resulting functional resilience of the ring is motif dependent. Defining these functional boundaries can help transform our understanding of AAA+ motors from structural snapshots into dynamic, coordinated, and adaptive molecular machines that balance mechanochemical efficiency with kinetic resilience.

## DATA AVAILABILITY

All processed data are available in the article and supporting information.

## ACKNOWLEDGMENTS

Funding for this work was provided by the Natural Sciences and Engineering Research Council of Canada (NSERC) Discovery Grants RGPIN-2020-00527 to R.H. We acknowledge the support from CFI- JEFL grants (43757 and 45118) to R.H. that provided the equipment access to this study.

## AUTHOR CONTRIBUTIONS

PK and RH conceptualized the project and designed experiments. PK performed all the experiments. PK and RH carried out data analysis and visualization. PK and RH wrote and edited the manuscript.

## DECLARATION OF INTERESTS

The authors declare no conflict of interest.

## Notes

### Competing Interest Statement

The authors have declared no competing interest.

## REFERENCES

(1) Khan, Y. A.; White, K. I.; Brunger, A. T. The AAA+ Superfamily: A Review of the Structural and Mechanistic Principles of These Molecular Machines. Crit. Rev. Biochem. Mol. Biol. 2022, 57 (2), 156–187. 10.1080/10409238.2021.1979460

(2) Puchades, C.; Sandate, C. R.; Lander, G. C. The Molecular Principles Governing the Activity and Functional Diversity of AAA+ Proteins. Nat. Rev. Mol. Cell Biol. 2020, 21 (1), 43–58. 10.1038/s41580-019-0183-6

(3) Carver, A.; Zhang, B.; Zhang, X. Structures and Mechanisms of AAA+ Protein Complexes in DNA Processing. Curr. Opin. Struct. Biol. 2025, 92, 103056. 10.1016/j.sbi.2025.103056

(4) Ripstein, Z. A.; Huang, R.; Augustyniak, R.; Kay, L. E.; Rubinstein, J. L. Structure of a AAA+ Unfoldase in the Process of Unfolding Substrate. Elife 2017, 6, e25754. 10.7554/eLife.25754

(5) Huang, R.; Ripstein, Z. A.; Augustyniak, R.; Lazniewski, M.; Ginalski, K.; Kay, L. E.; Rubinstein, J. L. Unfolding the Mechanism of the AAA+ Unfoldase VAT by a Combined Cryo-EM, Solution NMR Study. Proceedings of the National Academy of Sciences 2016, 113 (29), E4190–E4199. 10.1073/pnas.1603980113

(6) Puchades, C.; Rampello, A. J.; Shin, M.; Giuliano, C. J.; Wiseman, R. L.; Glynn, S. E.; Lander, G. C. Structure of the Mitochondrial Inner Membrane AAA+ Protease YME1 Gives Insight into Substrate Processing. Science (1979). 2017, 358 (6363), eaao0464. 10.1126/science.aao0464

(7) Puchades, C.; Ding, B.; Song, A.; Wiseman, R. L.; Lander, G. C.; Glynn, S. E. Unique Structural Features of the Mitochondrial AAA+ Protease AFG3L2 Reveal the Molecular Basis for Activity in Health and Disease. Mol. Cell 2019, 75 (5), 1073–1085.e6. 10.1016/j.molcel.2019.06.016

(8) de la Peña, A. H.; Goodall, E. A.; Gates, S. N.; Lander, G. C.; Martin, A. Substrate-Engaged 26S Proteasome Structures Reveal Mechanisms for ATP-Hydrolysis–Driven Translocation. Science (1979). 2018, 362 (6418), eaav0725. 10.1126/science.aav0725

(9) Gates, S. N.; Martin, A. Stairway to Translocation: AAA+ Motor Structures Reveal the Mechanisms of ATP-Dependent Substrate Translocation. Protein Science 2020, 29 (2), 407–419. 10.1002/pro.3743

(10) Lin, J.; Shorter, J.; Lucius, A. L. AAA+ Proteins: One Motor, Multiple Ways to Work. Biochem. Soc. Trans. 2022, 50 (2), 895–906. 10.1042/BST20200350

(11) Sauer, R. T.; Fei, X.; Bell, T. A.; Baker, T. A. Structure and Function of ClpXP, a AAA+ Proteolytic Machine Powered by Probabilistic ATP Hydrolysis. Crit. Rev. Biochem. Mol. Biol. 2022, 57 (2), 188–204. 10.1080/10409238.2021.1979461

(12) Martin, A.; Baker, T. A.; Sauer, R. T. Rebuilt AAA + Motors Reveal Operating Principles for ATP-Fuelled Machines. Nature 2005, 437 (7062), 1115–1120. 10.1038/nature04031

(13) Lin, J.; Banwait, J. K.; Saurabh, A.; Sharp, K. A.; Southworth, D. R.; Pressé, S.; Lucius, A. L.; Goldman, Y. E.; Shorter, J. Biased Stochastic Motor Dynamics Drive Bidirectional, Processive Translocation by the AAA+ Disaggregase Hsp104. bioRxiv 2026, 2026.08.09.743785. 10.64898/2026.08.09.743785

(14) DeSantis, M. E.; Leung, E. H.; Sweeny, E. A.; Jackrel, M. E.; Cushman-Nick, M.; Neuhaus- Follini, A.; Vashist, S.; Sochor, M. A.; Knight, M. N.; Shorter, J. Operational Plasticity Enables Hsp104 to Disaggregate Diverse Amyloid and Nonamyloid Clients. Cell 2012, 151 (4), 778–793. 10.1016/j.cell.2012.09.038

(15) Banwait, J. K.; Islam, L.; Lucius, A. L. Single Turnover Transient State Kinetics Reveals Processive Protein Unfolding Catalyzed by Escherichia Coli ClpB. Elife 2024, 13, RP99052. 10.7554/eLife.99052

(16) Casier, R.; Levy, D.; Riven, I.; Barak, Y.; Haran, G. A Stochastic Mechanism Drives Fast Substrate Translocation in the AAA+ Machine ClpB. Nat. Commun. 2026, 17 (1), 1773. 10.1038/s41467-026-68478-1

(17) Cordova, J. C.; Olivares, A. O.; Shin, Y.; Stinson, B. M.; Calmat, S.; Schmitz, K. R.; Aubin-Tam, M.-E.; Baker, T. A.; Lang, M. J.; Sauer, R. T. Stochastic but Highly Coordinated Protein Unfolding and Translocation by the ClpXP Proteolytic Machine. Cell 2014, 158 (3), 647–658. 10.1016/j.cell.2014.05.043

(18) Stinson, B. M.; Baytshtok, V.; Schmitz, K. R.; Baker, T. A.; Sauer, R. T. Subunit Asymmetry and Roles of Conformational Switching in the Hexameric AAA+ Ring of ClpX. Nat. Struct. Mol. Biol. 2015, 22 (5), 411–416. 10.1038/nsmb.3012

(19) Stinson, B. M.; Nager, A. R.; Glynn, S. E.; Schmitz, K. R.; Baker, T. A.; Sauer, R. T. Nucleotide Binding and Conformational Switching in the Hexameric Ring of a AAA+ Machine. Cell 2013, 153 (3), 628–639. 10.1016/j.cell.2013.03.029

(20) Gerega, A.; Rockel, B.; Peters, J.; Tamura, T.; Baumeister, W.; Zwickl, P. VAT, the Thermoplasma Homolog of Mammalian P97/VCP, Is an N Domain-Regulated Protein Unfoldase. Journal of Biological Chemistry 2005, 280 (52), 42856–42862. 10.1074/jbc.M510592200

(21) Werbeck, N. D.; Schlee, S.; Reinstein, J. Coupling and Dynamics of Subunits in the Hexameric AAA+ Chaperone ClpB. J. Mol. Biol. 2008, 378 (1), 178–190. 10.1016/j.jmb.2008.02.026

(22) Huang, R.; Ripstein, Z. A.; Rubinstein, J. L.; Kay, L. E. Cooperative Subunit Dynamics Modulate P97 Function. Proceedings of the National Academy of Sciences 2019, 116 (1), 158–167. 10.1073/pnas.1815495116

(23) Moreau, M. J.; McGeoch, A. T.; Lowe, A. R.; Itzhaki, L. S.; Bell, S. D. ATPase Site Architecture and Helicase Mechanism of an Archaeal MCM. Mol. Cell 2007, 28 (2), 304–314. 10.1016/j.molcel.2007.08.013

(24) Martin, A.; Baker, T. A.; Sauer, R. T. Pore Loops of the AAA+ ClpX Machine Grip Substrates to Drive Translocation and Unfolding. Nat. Struct. Mol. Biol. 2008, 15 (11), 1147–1151. 10.1038/nsmb.1503

(25) Iosefson, O.; Nager, A. R.; Baker, T. A.; Sauer, R. T. Coordinated Gripping of Substrate by Subunits of a AAA+ Proteolytic Machine. Nat. Chem. Biol. 2015, 11 (3), 201–206. 10.1038/nchembio.1732

