## Supplementary Material for "How Much Coordination Does an AAA+ Motor Require? A Case Study of the Archaeal Cdc48/p97 Homologue"

### Protein Sequences

#### ΔN-VAT (WT)

62969.39 Da

Predicted pI: 5.25

GEEVSRISYEDIGGLSEQLGKIREMIELPLKHPELFERLGITPPKGVILYGPPGTGKTLIARAVANE
SGANFLSINGPEIMSKYYGQSEQKLREIFSKAEETAPSIIFIDEIDSIAPKREEVQGEVERRVVAQL
LTLMDGMKERGHVIVIGATNRIDAIDPALRRPGRFDREIEIGVPDRNGRKEILMIHTRNMPLGMS
EEEKNKFLEEMADYTYGFVGADLAALVRESAMNALRRYLPEIDLDKPIPTEILEKMVVTEDDFKN
ALKSIEPSSLREVMVEVPNVHWDDIGGLEDVKREIKETVELPLLKPDVFKRLGIRPSKGFLLYGP
PGVGKTLLAKAVATESNANFISIKGPEVLSKWVGESEKAIREIFKKAKQVAPAIVFLDEIDSIAPRR
GTTSDSGVTERIVNQLLTSLDGIEVMNGVVVIGATNRPDIMDPALLRAGRFDKLIYIPPPDKEARL
SILKVHTKNMPLAPDVDLNDIAQRTEGYVGADLENLCREAGMNAYRENPDATSVSQKNFLDAL
KTIRPSVDEEVIKFYRTLSETMSKSVSERRKQLQDQGLYL

#### Full-length VAT (FL)

83195.81 Da

Predicted pI: 5.59

GMESNNGIILRVAEANSTDPGMSRVRLDESSRRLLDAEIGDVVEIEKVRKTVGRVYRARPEDEN
KGIVRIDSVMRNNCGASIGDKVKVRKVRTEIAKKVTLAPIIRKDQRLKFGEGIEEYVQRALIRRPM
LEQDNISVPGLTLAGQTGLLFKVVKTLPSKVPVEIGEETKIEIREEPASEVLEEVSRISYEDIGGLS
EQLGKIREMIELPLKHPELFERLGITPPKGVILYGPPGTGKTLIARAVANESGANFLSINGPEIMSK
YYGQSEQKLREIFSKAEETAPSIIFIDEIDSIAPKREEVQGEVERRVVAQLLTLMDGMKERGHVIV
IGATNRIDAIDPALRRPGRFDREIEIGVPDRNGRKEILMIHTRNMPLGMSEEEKNKFLEEMADYT
YGFVGADLAALVRESAMNALRRYLPEIDLDKPIPTEILEKMVVTEDDFKNALKSIEPSSLREVMV
EVPNVHWDDIGGLEDVKREIKETVELPLLKPDVFKRLGIRPSKGFLLYGPPGVGKTLLAKAVATE
SNANFISIKGPEVLSKWVGESEKAIREIFKKAKQVAPAIVFLDEIDSIAPRRGTTSDSGVTERIVNQ
LLTSLDGIEVMNGVVVIGATNRPDIMDPALLRAGRFDKLIYIPPPDKEARLSILKVHTKNMPLAPD
VDLNDIAQRTEGYVGADLENLCREAGMNAYRENPDATSVSQKNFLDALKTIRPSVDEEVIKFYR
TLSETMSKSVSERRKQLQDQGLYL

#### ΔN-VAT Walker A variant ($WA_v$)

62855.20 Da

Predicted pI: 5.16

GEEVSRISYEDIGGLSEQLGKIREMIELPLKHPELFERLGITPPKGVILYGPPGTG**<u>A</u>**TLIARAVAN
ESGANFLSINGPEIMSKYYGQSEQKLREIFSKAEETAPSIIFIDEIDSIAPKREEVQGEVERRVVA
QLLTLMDGMKERGHVIVIGATNRIDAIDPALRRPGRFDREIEIGVPDRNGRKEILMIHTRNMPLG
MSEEEKNKFLEEMADYTYGFVGADLAALVRESAMNALRRYLPEIDLDKPIPTEILEKMVVTEDD
FKNALKSIEPSSLREVMVEVPNVHWDDIGGLEDVKREIKETVELPLLKPDVFKRLGIRPSKGFLL
YGPPGVG**<u>A</u>**TLLAKAVATESNANFISIKGPEVLSKWVGESEKAIREIFKKAKQVAPAIVFLDEIDSIA
PRRGTTSDSGVTERIVNQLLTSLDGIEVMNGVVVIGATNRPDIMDPALLRAGRFDKLIYIPPPDK

EARLSILKVHTKNMPLAPDVDLNDIAQRTEGYVGADLENLCREAGMNAYRENPDATSVSQKNF
LDALKTIRPSVDEEVIKFYRTLSETMSKSVSERRKQLQDQGLYL

**ΔN-VAT Arginine Finger variant (AF$_v$)**

62799.17 Da

Predicted pI: 5.16

GEEVSRISYEDIGGLSEQLGKIREMIELPLKHPELFERLGITPPKGVILYGPPGTGKTLIARAVANE
SGANFLSINGPEIMSKYYGQSEQKLREIFSKAEETAPSIIFIDEIDSIAPKREEVQGEVERRVVAQL
LTLMDGMKERGHVIVIGATNRIDAIDPALR**A**PGRFDREIEIGVPDRNGRKEILMIHTRNMPLGMS
EEEKNKFLEEMADYTYGFVGADLAALVRESAMNALRRYLPEIDLDKPIPTEILEKMVVTEDDFKN
ALKSIEPSSLREVMVEVPNVHWDDIGGLEDVKREIKETVELPLLKPDVFKRLGIRPSKGFLLYGP
PGVGKTLLAKAVATESNANFISIKGPEVLSKWVGESEKAIREIFKKAKQVAPAIVFLDEIDSIAPRR
GTTSDSGVTERIVNQLLTSLDGIEVMNGVVVIGATNRPDIMDPALL**A**AGRFDKLIYIPPPDKEARL
SILKVHTKNMPLAPDVDLNDIAQRTEGYVGADLENLCREAGMNAYRENPDATSVSQKNFLDAL
KTIRPSVDEEVIKFYRTLSETMSKSVSERRKQLQDQGLYL

**ΔN-VAT Pore Loop variant (PL$_v$)**

62642.01 Da

Predicted pI: 5.25

GEEVSRISYEDIGGLSEQLGKIREMIELPLKHPELFERLGITPPKGVILYGPPGTGKTLIARAVANE
SGANFLSINGPEIMSK**AA**GQSEQKLREIFSKAEETAPSIIFIDEIDSIAPKREEVQGEVERRVVAQ
LLTLMDGMKERGHVIVIGATNRIDAIDPALRRPGRFDREIEIGVPDRNGRKEILMIHTRNMPLGM
SEEEKNKFLEEMADYTYGFVGADLAALVRESAMNALRRYLPEIDLDKPIPTEILEKMVVTEDDFK
NALKSIEPSSLREVMVEVPNVHWDDIGGLEDVKREIKETVELPLLKPDVFKRLGIRPSKGFLLYG
PPGVGKTLLAKAVATESNANFISIKGPEVLSK**AA**GESEKAIREIFKKAKQVAPAIVFLDEIDSIAPR
RGTTSDSGVTERIVNQLLTSLDGIEVMNGVVVIGATNRPDIMDPALLRAGRFDKLIYIPPPDKEA
RLSILKVHTKNMPLAPDVDLNDIAQRTEGYVGADLENLCREAGMNAYRENPDATSVSQKNFLD
ALKTIRPSVDEEVIKFYRTLSETMSKSVSERRKQLQDQGLYL

### Supplementary Tables

**Table S1. Comparison of the fitting models for NADH-coupled ATPase assay.**

| | Unidirectional | Bidirectional | Independent | All-or-None |
|---|---|---|---|---|
| **WT + $WA_v$** | | | | |
| $\chi^2$ | 3.57 | 3.01 | 112.97 | 1247.68 |
| **df** | 5 | 5 | 6 | |
| $\chi_r^2$ | 0.71 | 0.60 | 18.83 | 207.95 |
| $\Delta AIC_c$ | 0.56 | 0.00 | 106.96 | 1241.67 |
| $\alpha \pm SE$ | -0.62 ± 0.05 | | | |
| $\alpha' \pm SE$ | | -0.33 ± 0.03 | | |
| **95% CI** ($\alpha$ **or** $\alpha'$) | -0.51, -0.74 | -0.27, -0.40 | | |
| **WT + $AF_v$** | | | | |
| $\chi^2$ | 2.40 | 6.81 | 1367.00 | 578.00 |
| **df** | 5 | 5 | 6 | 6 |
| $\chi_r^2$ | 0.48 | 1.36 | 227.83 | 96.33 |
| $\Delta AIC_c$ | 0.00 | 4.41 | 1361.60 | 572.60 |
| $\alpha \pm SE$ | -0.97 ± 0.02 | | | |
| $\alpha' \pm SE$ | | -0.60 ± 0.03 | | |
| **95% CI** ($\alpha$ **or** $\alpha'$) | -0.92, -1.00 | -0.56, -0.65 | | |
| **WT + $PL_v$** | | | | |
| $\chi^2$ | 1.06 | 1.60 | 38.56 | 74.97 |
| **df** | 5 | 5 | 6 | 6 |
| $\chi_r^2$ | 0.21 | 0.32 | 6.43 | 12.49 |
| $\Delta AIC_c$ | 0.00 | 0.54 | 34.50 | 70.91 |
| $\alpha \pm SE$ | +1.29 ± 0.10 | | | |
| $\alpha' \pm SE$ | | +0.55 ± 0.00 | | |
| **95% CI** ($\alpha$ **or** $\alpha'$) | +0.88, +1.40 | +0.40, +0.55 | | |
| **WT + FL** | | | | |
| $\chi^2$ | 3.38 | 3.38 | 3.39 | 130.57 |
| **df** | 5 | 5 | 6 | 6 |
| $\chi_r^2$ | 0.68 | 0.68 | 0.56 | 21.76 |
| $\Delta AIC_c$ | 2.99 | 2.99 | 0.00 | 127.19 |
| $\alpha \pm SE$ | -0.02 ± 0.13 | | | |
| $\alpha' \pm SE$ | | -0.01 ± 0.07 | | |
| **95% CI** ($\alpha$ **or** $\alpha'$) | 0.00, -0.32 | 0.00, -0.17 | | |

* df: degree of freedom; SE: standard error from the covariance matrix; CI: confidence interval.

**Table S2. Comparison of the fitting models for substrate unfolding assay.**

| | Threshold + Exponential (T-E) | Threshold | Exponential | Independent | All-or-None |
|---|---|---|---|---|---|
| **WT + WA$_v$** | | | | | |
| $\chi^2$ | 1.72 | 21.65 | 18.83 | 57.25 | 187.99 |
| **df** | 4 | 5 | 5 | 6 | 6 |
| $\chi_r^2$ | 0.43 | 4.33 | 3.77 | 9.54 | 31.33 |
| $\Delta\mathbf{AIC_c}$ | 0.00 | 14.93 | 12.11 | 47.53 | 178.27 |
| $\boldsymbol{T}$ | 4 | 3 | | | |
| $\boldsymbol{\beta} \pm \mathbf{SE}$ | 0.28 ± 0.02 | | 0.39 ± 0.05 | | |
| **95% CI ($\boldsymbol{\beta}$)** | 0.23, 0.34 | | 0.35, 0.43 | | |
| **WT + AF$_v$** | | | | | |
| $\chi^2$ | 1.39 | 27.75 | 68.59 | 33.04 | 747.47 |
| **df** | 4 | 5 | 5 | 6 | 6 |
| $\chi_r^2$ | 0.35 | 5.55 | 13.72 | 5.51 | 124.58 |
| $\Delta\mathbf{AIC_c}$ | 0.00 | 21.36 | 62.20 | 23.64 | 738.08 |
| $\boldsymbol{T}$ | 4 | 3 | | | |
| $\boldsymbol{\beta} \pm \mathbf{SE}$ | 0.18 ± 0.01 | | 0.31 ± 0.04 | | |
| **95% CI ($\boldsymbol{\beta}$)** | 0.15, 0.21 | | 0.30, 0.33 | | |
| **WT + PL$_v$** | | | | | |
| $\chi^2$ | 3.47 | 6.61 | 10.89 | 120.83 | 43.86 |
| **df** | 4 | 5 | 5 | 6 | 6 |
| $\chi_r^2$ | 0.87 | 1.32 | 2.18 | 20.14 | 7.31 |
| $\Delta\mathbf{AIC_c}$ | 1.86 | 0.00 | 4.28 | 111.22 | 34.25 |
| $\boldsymbol{T}$ | 3 | 2 | | | |
| $\boldsymbol{\beta} \pm \mathbf{SE}$ | 0.31 ± 0.07 | | 0.52 ± 0.07 | | |
| **95% CI ($\boldsymbol{\beta}$)** | 0.17, 0.47 | | 0.45, 0.62 | | |
| **WT + FL** | | | | | |
| $\chi^2$ | 3.32 | 36.13 | 20.85 | 2.97 | 201.42 |
| **df** | 4 | 5 | 5 | 6 | 6 |
| $\chi_r^2$ | 0.83 | 7.23 | 4.17 | 0.49 | 33.57 |
| $\Delta\mathbf{AIC_c}$ | 8.35 | 36.16 | 20.89 | 0.00 | 198.45 |
| $\boldsymbol{T}$ | 6 | 4 | | | |
| $\boldsymbol{\beta} \pm \mathbf{SE}$ | 0.24 ± 0.01 | | 0.30 ± 0.03 | | |
| **95% CI ($\boldsymbol{\beta}$)** | 0.21, 0.28 | | 0.27, 0.33 | | |

* df: degree of freedom; SE: standard error from the covariance matrix; CI: confidence interval.

### Supplementary Figures

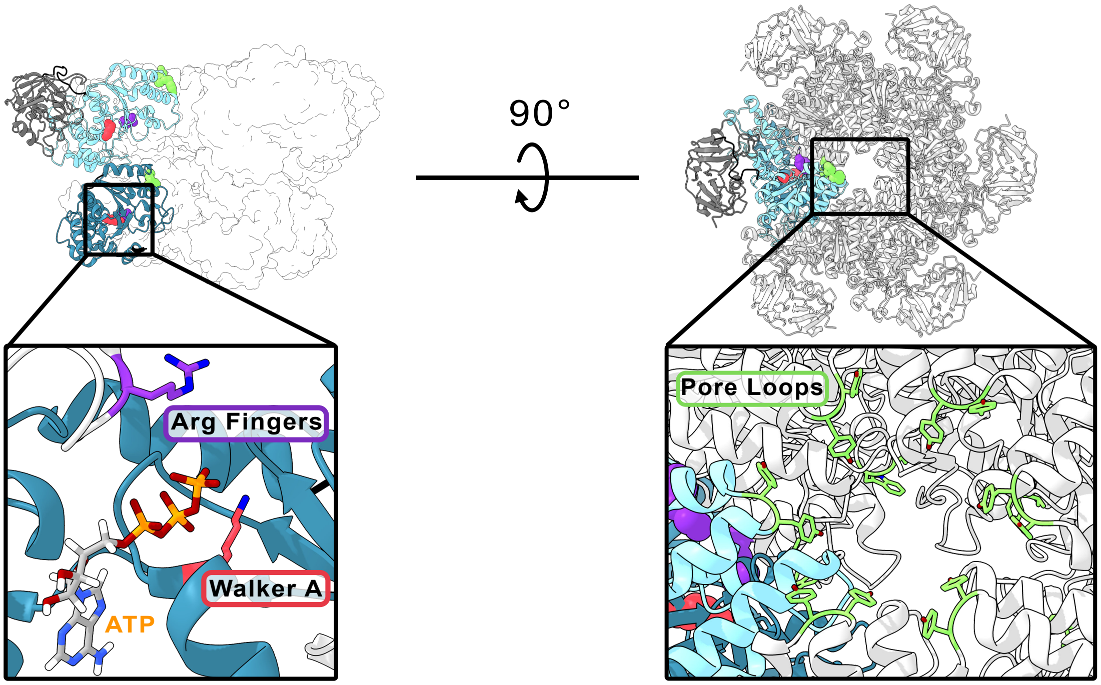


**Figure S1. Conserved mechanochemical motifs in full-length VAT.** Cryo-EM structure of full-length VAT in the symmetric, stacked-ring conformation (PDB: 5G4G).[1] Insets highlight in stick representation the conserved Walker A motif (red), Arginine finger (purple), and ATP in the nucleotide binding pocket (left), and the pore loops (green) lining the central pore (right).

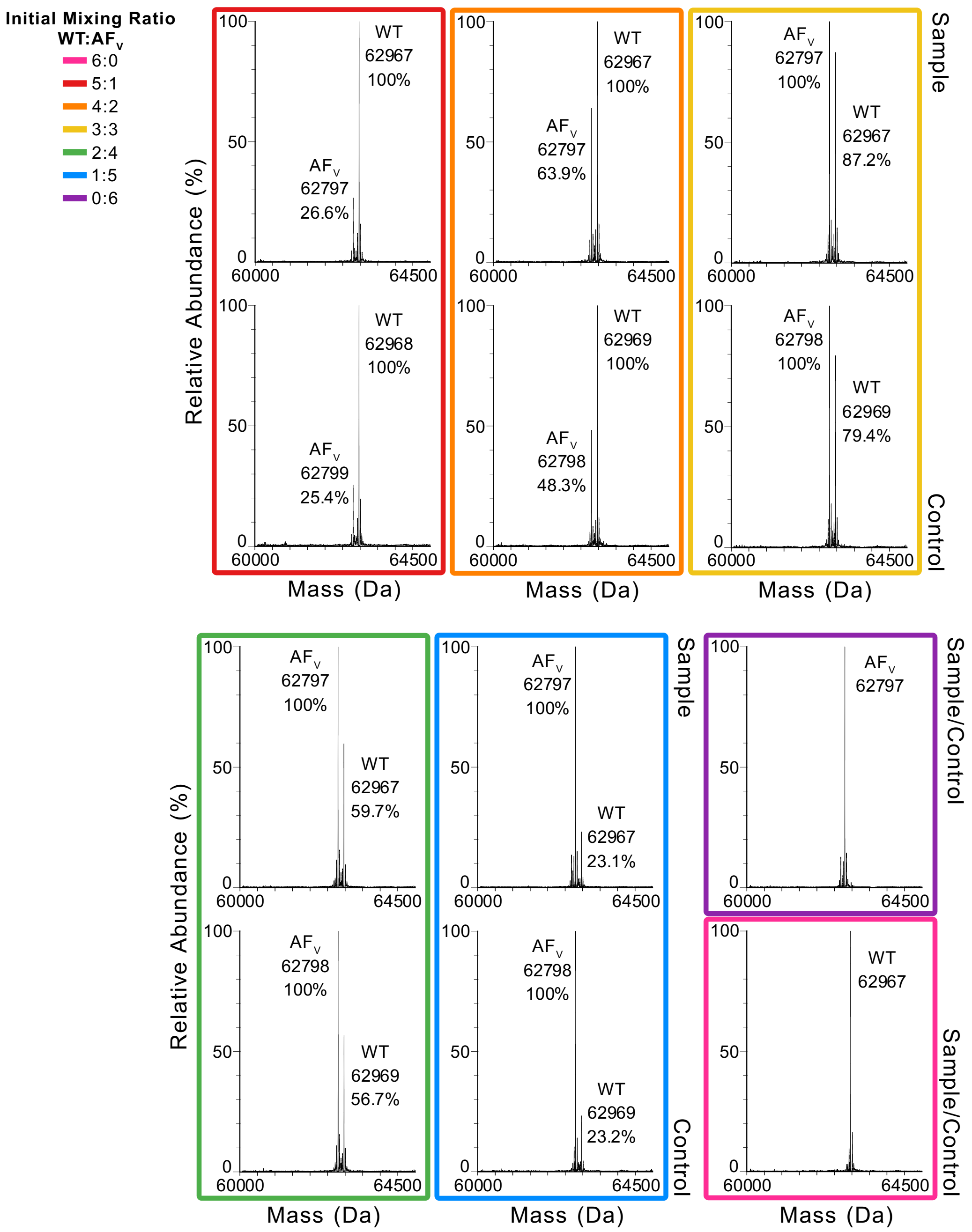


**Figure S2. Validation of protomer composition in reconstituted WT-AF$_V$ heterohexamers using ESI-MS**. The colored rectangles indicate the initial mixing ratios of WT and AF$_V$ protomers. ESI-MS spectra of the mixed samples were acquired from the mixed samples before unfolding–refolding (Control) and from the hexameric fractions collected after unfolding–refolding and size-exclusion chromatography (Sample; Figure 1E). The relative molar ratios of WT and AF$_V$ in the Sample were normalized to those in the corresponding Control to account for differences in ionization efficiencies between the two protomers.

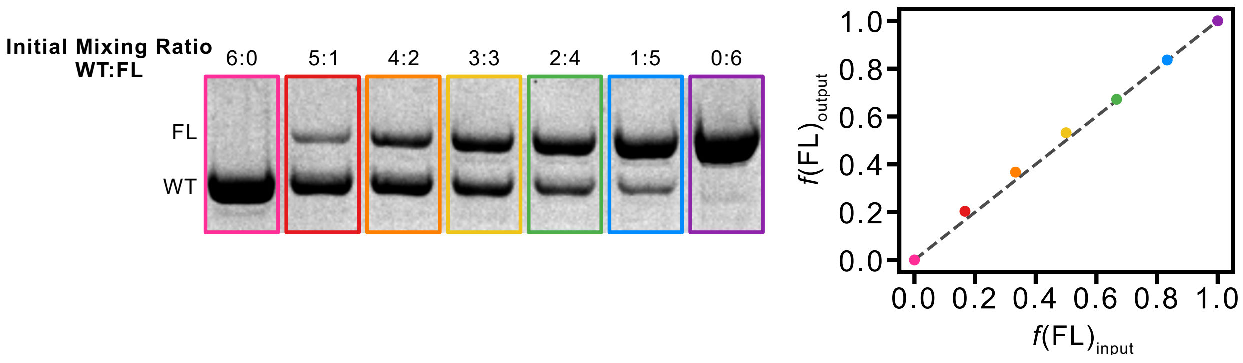


**Figure S3. Validation of protomer composition in reconstituted WT-FL heterohexamers by SDS-PAGE**. The colored rectangles indicate the initial mixing ratios of WT and FL protomers. Samples collected from the hexameric fractions following size-exclusion chromatography of the unfolding–refolding mixtures (Figure 1E) were analyzed by SDS-PAGE (left). The relative band intensities, corrected for the molecular weights of the two protomers, were used to determine their molar ratios in each sample. Corrected molar ratios of the refolded samples are plotted against the input mixing ratios (right). The grey dashed line indicates y=x.

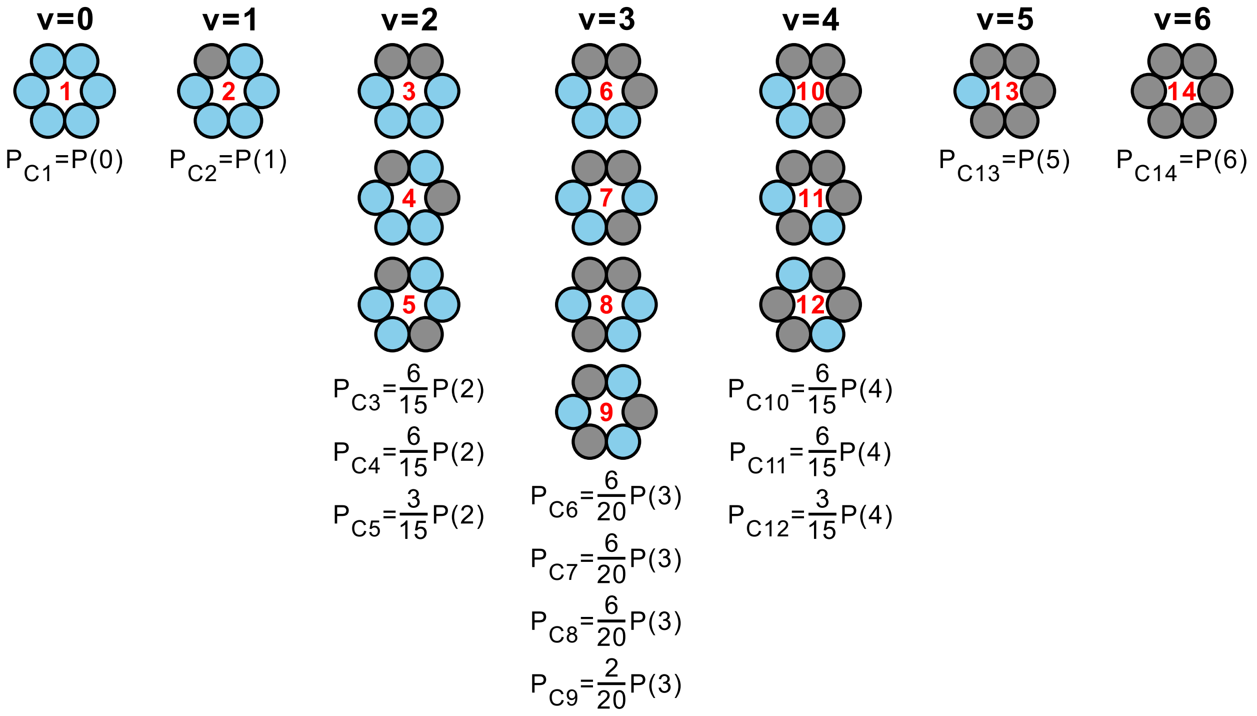


**Figure S4. Populations of distinct ring configurations in the reconstituted heterohexamer samples.** The 14 possible configurations are arranged in columns according to the number of variant protomers in the ring ($v$ = 0–6). The probability of each specific configuration ($P_C$) is calculated by weighting the binomial probability of its stoichiometric class, $P(v)$, by the fraction of distinct rotational arrangements represented by that configuration.

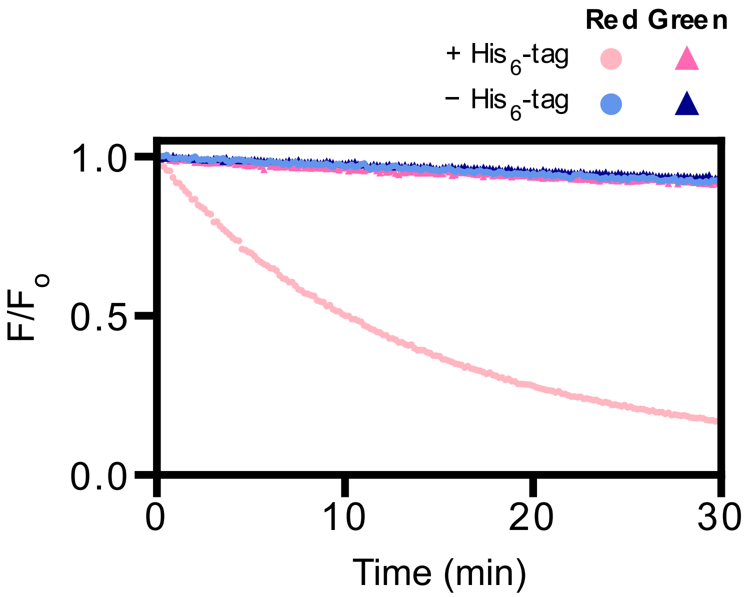


**Figure S5. VAT requires an unstructured terminus for unfolding.** The green (excitation/emission: 480/516 nm) and red (excitation/emission: 540/580 nm) fluorescence intensities of photoconverted mEos3.2-tagged substrate, with (+) or without (-) an unstructured N-terminal $His_6$-tag, were monitored over time and normalized to their respective initial values. The unfolding assay was performed with final protomer concentrations of 23 $\mu$M ΔN-WT VAT and 2.3 $\mu$M substrate in the presence of an ATP regeneration system.

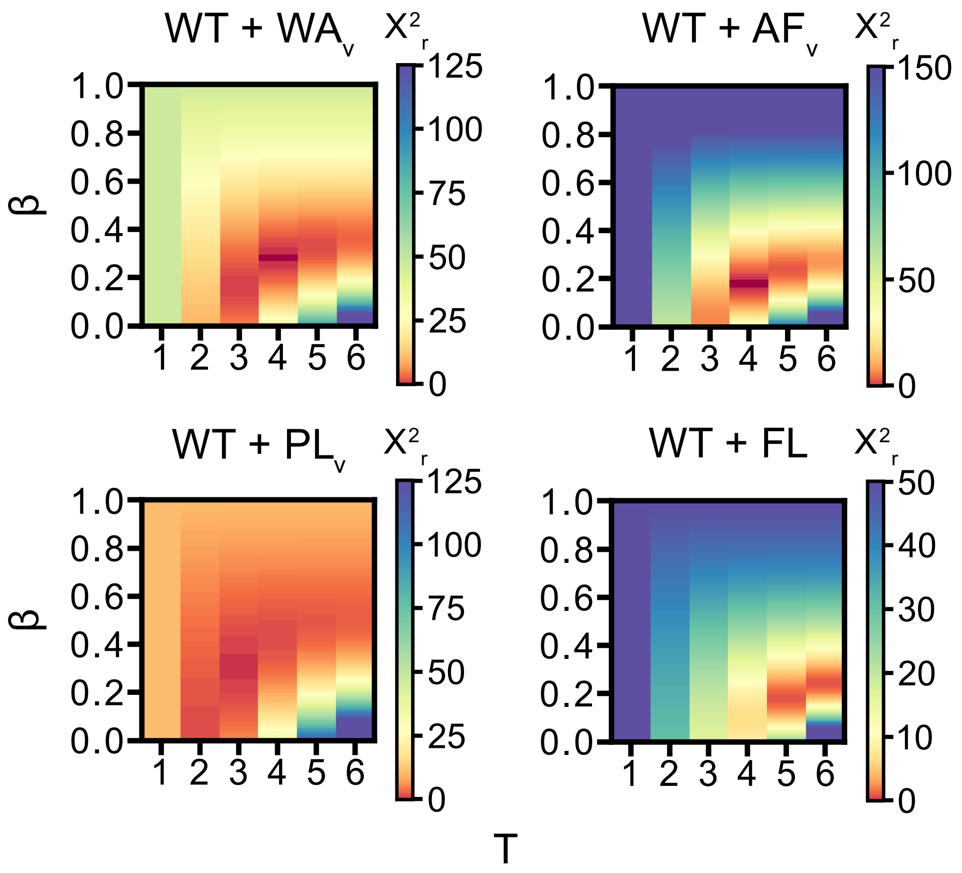


**Figure S6. Systematic grid search to determine optimal parameter set for Threshold+Exponential model.** The Threshold+Exponential model includes both threshold ($T$) and exponential inhibitory ($\beta$) parameters. To identify the optimal set, a systematic grid search of $\chi_r^2$ values was used, where minimum values indicate the best fit (red).

### Materials and Methods

#### Cloning, Protein Expression, and Purification

The *T. acidophilum* VAT protein was encoded in a pProEx-1 vector with an N-terminal $His_6$-tag and a tobacco etch virus (TEV) protease cleavage site. Additional constructs, including ΔN-VAT (WT) (residues 182-745) and the ΔN mutants $WA_v$ (K237A, K514A), $PL_v$ (Y264A, Y265A, W541A, V542A), and $AF_v$ (R345A, R623A), were generated via QuikChange site-directed mutagenesis using Phusion DNA polymerase (New England Biolabs) and primers produced by Integrated DNA Technologies (IDT). All plasmids were transformed into *E. coli* 10-beta cells, isolated using the GenElute plasmid miniprep kit (Sigma-Aldrich), and verified via full-plasmid sequencing (Plasmidsaurus).

All VAT expression vectors were transformed into *E. coli* BL21(DE3) cells and grown in LB broth supplemented with 100 mg/L ampicillin at 37 °C. Protein expression was induced with 0.25 mM isopropyl β-D-1-thiogalactopyranoside (IPTG) at an $OD_{600}$ of 0.6, followed by incubation for 16 hours at 25 °C.

Cells were harvested by centrifugation and resuspended in Ni-A buffer [50 mM tris (pH 7.4), 500 mM NaCl, 50 mM imidazole]. The cells were lysed with ~1 mg of lysozyme, followed by two 10 min cycles of sonication (45% amplitude, 2-s on/2-s off pulses) with mixing between cycles. The lysate was incubated in a 50 °C water bath for 10 mins and subsequently centrifuged at 8,000 rpm for 20 mins. The supernatant was loaded onto a column containing 5 mL of HisPur Ni-NTA resin (Thermo Scientific) previously equilibrated with Ni-A buffer. After washing the column with Ni-A buffer, the bound target protein was eluted with Ni-B buffer [25 mM tris (pH 7.8), 300 mM imidazole]. The $His_6$-tag was cleaved by adding TEV protease to the eluate (molar ratio TEV:protein = 1:50), followed by overnight dialysis at 20 °C against a buffer containing 50 mM tris (pH 7.4), 200 mM NaCl, 0.5 mM EDTA, 0.5 mM DTT. To separate the cleaved target protein from the $His_6$-tag, uncleaved protein, and TEV protease, the dialyzed sample was subjected to a second round of Ni-NTA chromatography as described above. The flowthrough containing the cleaved target protein was collected and concentrated using an Amicon Ultra centrifugal filter (MilliporeSigma). This concentrated sample was further purified via size exclusion chromatography (SEC) by loading it onto a Superdex 200 Increase 10/300 GL column (Cytiva) equilibrated and run in SEC buffer [50 mM HEPES (pH 7.5), 100 mM NaCl, 1 mM EDTA].

Protein purity was assessed by SDS-PAGE and visualized using Image Lab (Bio-Rad). Fractions containing pure protein were pooled, and the protein concentration was calculated via the Beer-Lambert law using the measured absorbance at 280 nm and extinction coefficients derived from the ExPASy ProtParam tool. The pure protein was flash-frozen in liquid nitrogen and stored at -80 °C.

#### Population Distribution of Ring Configurations

Reconstituted mixed heterohexamers were prepared by initially mixing WT and variant homohexamers at defined molar ratios. The fraction of variant protomers in the mixture, $f_{\mathrm{Var}}$, is defined as

$$f_{\mathrm{Var}} = \frac{[\mathrm{Var}]}{[\mathrm{Var}]+[\mathrm{WT}]} \qquad \text{Eq. (1)}$$

where $[\mathrm{Var}]$ and $[\mathrm{WT}]$ denotes the concentrations of variant and WT protomers, respectively. Assuming random protomer assembly during unfolding–refolding,[2,3] the resulting sample contains 14 distinct ring configurations with their populations following a binomial distribution determined by $f_{\mathrm{Var}}$ (Figure 1C,D).[1,4] The probability, $P(v)$, of hexameric rings containing *v* variant protomers is given by

$$P(v) = \frac{6!}{v!\,(6-v)!} {f_{\mathrm{Var}}}^{v} (1 - f_{\mathrm{Var}})^{6-v}$$

The fractional population of an individual ring configuration, $P_{\mathrm{C}}$, is then obtained by weighting $P(v)$ by the fraction of distinct spatial arrangements within the corresponding stoichiometric class represented by that configuration (Figure S4).

#### Models for ATPase Activity

For both ATPase and substrate unfolding activities, the experimentally observed activity, $A(f_{\mathrm{Var}})$, of a mixed heterohexamer sample is calculated as the population-weighted average of the activities of all possible ring configurations

$$A(f_{\mathrm{Var}}) = \sum_{C=1}^{14} P_C \times A_C$$

where $f_{\mathrm{Var}}$ is the fraction of variant protomers in the mixture, $P_C$ is fractional population of ring configuration *C*, and $A_C$ is the ATPase activity of a hexamer with that configuration.

*Independent Model*. In the Independent model, the ATPase activity of a hexameric ring is the weighted sum of the activities of its WT and variant protomers:

$$A_{\mathrm{c}} = f_{\mathrm{WT}} \cdot A_{\mathrm{WT}} + f_{\mathrm{Var}} \cdot A_{\mathrm{Var}}$$

where $f_{\mathrm{WT}}$ and $f_{\mathrm{Var}}$ are the fractions of the WT and variant protomers, respectively, in a given ring configuration, and $A_{\mathrm{WT}}$ and $A_{\mathrm{Var}}$ are the ATPase activities of the corresponding WT and variant homohexamers. This model assumes that promoters act independent without inter-protomer coupling.

*All-or-None Model*. In the All-or-None model, incorporation of one or more variant protomers shifts the activity of the entire hexamer to that of the variant homohexamers. The activity of a ring containing $v$ variant protomers is given by

$$A_{\mathrm{c}} = A(v) = \begin{cases} A_{\mathrm{WT}}, & v = 0 \\ A_{\mathrm{Var}}, & v \geq 1 \end{cases}$$

where $v$ is the number of variant protomers in the ring, and $A_{\mathrm{WT}}$ and $A_{\mathrm{Var}}$ are the activities of the WT and variant homohexamers, respectively.

*Unidirectional and Bidirectional Models.* In these models, the ATPase activity of a hexameric ring, $A_{\mathrm{c}}$, is calculated as the mean activity of its six protomers.

$$A_C = \frac{1}{6}\sum_{i=1}^{6} A_i$$

where $A_i$ is the ATPase activity of a given protomer $i$ according to its identity and the identities of its neighboring protomers.

In the Unidirectional model, the activity of a protomer $i$ depends on its own identity and the identity of its immediately preceding neighbor in the counterclockwise direction.

$$A_i = \begin{cases} A_{\mathrm{WT}}, & i = \mathrm{WT},\ i\text{-}1 = \mathrm{WT} \\ (1+\alpha)A_{\mathrm{WT}}, & i = \mathrm{WT},\ i\text{-}1 = \mathrm{Var} \\ A_{\mathrm{Var}}, & i = \mathrm{Var} \end{cases}$$

where $\alpha$ is the coupling parameter describing the effect of a neighboring variant protomer on the activity of a WT protomer. When $\alpha < 0$, the neighboring variant protomer inhibits the activity of the WT protomer; when $\alpha = 0$, it has no effect; and when $\alpha > 0$, it stimulates WT protomer activity. The value of $\alpha$ ranges from -1 (complete inhibition) to $A_{\mathrm{PL_v}}/A_{\mathrm{WT}}$ -1 (maximum activation).

In the Bidirectional model, the ATPase activity of protomer $i$ depends on its own identity and the identities of both adjacent neighbors:

$$A_i = \begin{cases} A_{\mathrm{WT}}, & i = \mathrm{WT},\ i\text{-}1 \text{ and } i\text{+}1 = \mathrm{WT} \\ (1+\alpha')A_{\mathrm{WT}}, & i = \mathrm{WT},\ i\text{-}1 \text{ or } i\text{+}1 = \mathrm{Var} \\ (1+\alpha')^2 A_{\mathrm{WT}}, & i = \mathrm{WT},\ i\text{-}1 \text{ and } i\text{+}1 = \mathrm{Var} \\ A_{\mathrm{Var}}, & i = \mathrm{Var} \end{cases}$$

where the coupling parameter $\alpha'$ describes the effect of the adjacent variant protomers on WT protomer activity. As in the Unidirectional model, $\alpha' < 0$ indicates inhibition by neighboring variant protomers, $\alpha' = 0$ indicates no effect, and $\alpha' > 0$ indicates activation. When both adjacent neighbors are variant protomers, their effects are assumed to be multiplicative. The value of $\alpha'$ ranges from -1 (complete inhibition) to $\sqrt{A_{\mathrm{PL_v}}/A_{\mathrm{WT}}}$ -1 (maximum activation).

#### Models for Unfoldase Activity

*Independent Model*. In the Independent model, the substrate unfolding activity of a hexameric ring is the weighted sum of the activities of its WT and variant protomers:

$$A_{\mathrm{c}} = f_{\mathrm{WT}} \cdot A_{\mathrm{WT}} + f_{\mathrm{Var}} \cdot A_{\mathrm{Var}}$$

where $f_{\mathrm{WT}}$ and $f_{\mathrm{Var}}$ are the fractions of the WT and variant protomers, respectively, in a given ring configuration, and $A_{\mathrm{WT}}$ and $A_{\mathrm{Var}}$ are the unfolding activities of the corresponding WT and variant homohexamers. This model assumes that each protomer contribute independently to substrate unfolding without inter-protomer coupling.

*All-or-None Model*. In the All-or-None model, incorporation of one or more variant protomers shifts the activity of the entire hexamer to that of the variant homohexamers. The activity of a ring containing $v$ variant protomers is given by

$$A_{\mathrm{c}} = A(v) = \begin{cases} A_{\mathrm{WT}}, & v = 0 \\ A_{\mathrm{Var}}, & v \geq 1 \end{cases}$$

where $v$ is the number of variant protomers in the ring, and $A_{\mathrm{WT}}$ and $A_{\mathrm{Var}}$ are the unfolding activities of the WT and variant homohexamers, respectively.

*Threshold Model.* In the Threshold model, the hexameric ring retains WT-level unfolding activity until the number of variant protomers reaches a threshold, $T$, beyond which it shifts to the variant homohexamer level. The activity of a ring containing $v$ variant protomers is given by

$$A_{\mathrm{c}} = A(v) = \begin{cases} A_{\mathrm{WT}}, & v < T \\ A_{\mathrm{Var}}, & v \geq T \end{cases}$$

where $v$ is the number of variant protomers in the ring, $T$ is the threshold, and $A_{\mathrm{WT}}$ and $A_{\mathrm{Var}}$ are the unfolding activities of the WT and variant homohexamers, respectively.

*Exponential Model.* In the Exponential model, the unfolding activity of a hexamer decreases exponentially from the WT toward the variant homohexamer level as the number of variant protomers increases. The activity of a ring containing $v$ variant protomers, $A(v)$, is given by

$$A_{\mathrm{c}} = A(v) = A_{\mathrm{Var}} + (A_{\mathrm{WT}} - A_{\mathrm{Var}})(1 - \beta)^{v}$$

where $\beta$ is the inhibitory parameter ranging from 0 (no inhibition) to 1 (complete inhibition), and $A_{\mathrm{WT}}$ and $A_{\mathrm{Var}}$ are the unfolding activities of the WT and variant homohexamers, respectively.

*Threshold+Exponential (T-E) Model.* The Threshold+Exponential (T-E) model combines the features of the Threshold and Exponential models. The unfolding activity of a hexameric ring decreases exponentially as the number of variant protomers increases, until the number reaches the threshold, $T$, at which point the activity shifts to the level of the variant homohexamers, described below:

$$A_{\mathrm{c}} = A(v) = \begin{cases} A_{\mathrm{Var}} + (A_{\mathrm{WT}} - A_{\mathrm{Var}})(1 - \beta)^{v}, & v < T \\ A_{\mathrm{Var}}, & v \geq T \end{cases}$$

where $v$ is the number of variant protomers, $T$ is the threshold number of variant protomers, and $\beta$ is the inhibitory parameter ranging from 0 (no inhibition) to 1 (complete inhibition). $A_{\mathrm{WT}}$ and $A_{\mathrm{Var}}$ are the unfolding activities of the WT and variant homohexamers, respectively.

**Evaluation of Goodness-of-Fit and Model Comparison**

The goodness-of-fit of each proposed model was evaluated using reduced chi-square ($\chi_r^2$), calculated by normalizing chi-square ($\chi^2$) by the degree of freedom

$$\chi^2 = \sum_{i=1}^{N} \left( \frac{A(f_{\mathrm{Var}_i}) - M(f_{\mathrm{Var}_i})}{\sigma\left(A(f_{\mathrm{Var}_i})\right)} \right)^2, \; \chi_r^2 = \frac{\chi^2}{N-p}$$

where $A(f_{\mathrm{Var}_i})$, $M(f_{\mathrm{Var}_i})$, $\sigma\left(A(f_{\mathrm{Var}_i})\right)$ are the experimental activity, model-predicted activity, and uncertainty of the experimental activity at a specific variant fraction $\left(f_{\mathrm{Var}_i}\right)$ calculated from experimental replicates. $N$ is the number of data points and $p$ is the number of fitting parameters in the model. A reduced chi-square value ($\chi_r^2$) close to 1 indicates that the model fits the data within the expected experimental uncertainty. Values substantially greater than 1 suggest a poorer fit relative to the assumed experimental uncertainty, whereas values substantially below 1 may indicate overfitting or an overestimation of experimental uncertainty.[5]

To compare candidate models while accounting for differences in model complexity, the corrected Akaike Information Criterion ($\mathrm{AIC_C}$) was calculated for each model, given by

$$AIC_C = \chi^2 + 2p + \frac{2p(p+1)}{N-p-1}$$

where $N$ is the number of data points and $p$ is the number of fitting parameters in the model. The relative support for each model was assessed using the difference between its AICc value and the minimum AICc among all candidate models, $\Delta AICc_i$:

$$\Delta AICc_i = AICc_i - AICc_{min}$$

where $AICc_i$ is the $AICc$ value of model $i$, and $AICc_{min}$ is the lowest $AICc$ value among the models considered. Models with $\Delta AICc_i \leq 2$ are generally considered to have substantial support relative to the best-supported model, whereas models with $\Delta AICc_i > 10$ indicates considerably less support.

### References


(1) Ripstein, Z. A.; Huang, R.; Augustyniak, R.; Kay, L. E.; Rubinstein, J. L. Structure of a AAA+ Unfoldase in the Process of Unfolding Substrate. *Elife* 2017, *6*, e25754. https://doi.org/10.7554/eLife.25754

(2) DeSantis, M. E.; Leung, E. H.; Sweeny, E. A.; Jackrel, M. E.; Cushman-Nick, M.; Neuhaus-Follini, A.; Vashist, S.; Sochor, M. A.; Knight, M. N.; Shorter, J. Operational Plasticity Enables Hsp104 to Disaggregate Diverse Amyloid and Nonamyloid Clients. *Cell* 2012, *151* (4), 778–793. https://doi.org/10.1016/j.cell.2012.09.038

(3) Huang, R.; Ripstein, Z. A.; Rubinstein, J. L.; Kay, L. E. Cooperative Subunit Dynamics Modulate P97 Function. *Proceedings of the National Academy of Sciences* 2019, *116* (1), 158–167. https://doi.org/10.1073/pnas.1815495116

(4) Werbeck, N. D.; Schlee, S.; Reinstein, J. Coupling and Dynamics of Subunits in the Hexameric AAA+ Chaperone ClpB. *J. Mol. Biol.* 2008, *378* (1), 178–190. https://doi.org/https://doi.org/10.1016/j.jmb.2008.02.026

(5) Burnham, K. P.; Anderson, D. R. *Model Selection and Multi-Model Inference : A Practical Information-Theoretic Approach*, 2nd ed.; Springer: Secaucus, UNITED STATES, 2002.
